# An update on the model of connectivity of the hippocampal formation (I): the perforant pathway to the dentate gyrus

**DOI:** 10.64898/2026.04.23.720458

**Authors:** Jon I Arellano, Pasko Rakic

## Abstract

The hippocampus participates in crucial functions such as memory consolidation, spatial processing and emotional regulation that require diverse input from multiple cortical areas that is funneled through the upper layers of the entorhinal cortex (EC), mostly from layer II to the dentate gyrus (DG). Traditional models described 200,000 EC layer II neurons projecting to 1 million granule cells (GCs) in the rat, rendering low divergence (1:5), with each EC neuron establishing about 18,000 synapses with GCs and each GC receiving about 4,000 synapses from EC neurons. In this manuscript, we update this model of connectivity incorporating new features described in the last three decades that include updated populations of EC layer II neurons obtained with design-based stereology, a revised definition of EC layer II based on molecular criteria and selecting reelin expressing neurons as the only layer II neurons projecting to the hippocampus. The updated model shows ∼80,000 neurons from EC layer II projecting to the DG, ∼45,000 from the medial entorhinal cortex (MEC) and ∼35,000 from the lateral entorhinal cortex (LEC) with high divergence of 1:20 and 1:30. We also show that EC layer II neurons may establish ∼90,000-115,000 synapses on GCs, while GCs receive about 8,000 synapses from EC layer II neurons. We estimate a ∼25% redundancy in the connectivity, so each EC neuron may contact ∼68,000-86,000 GCs and each GC would be contacted by ∼3000 neurons from MEC and 3,000 from LEC. In addition, we quantitatively assess a potential projection of mossy cells to the medial molecular layer described in mice, which could have a potential impact on GC inhibition. Overall, we produced a detailed, complete, and updated quantitative model of EC projections to the DG that reveals a much more divergent and richer projection than previously described, with implications for functional models (e.g.: pattern separation) and more widely for building realistic hippocampal models or establishing comparisons across species.

## Introduction

The hippocampal formation has a relevant role in cognitive functions such as memory consolidation, spatial processing, and emotional regulation. Its well-defined cytoarchitecture and spatially segregated, mostly unidirectional connections, has made it a preferred subject for structural and functional modelling in mammals. Rodents have been the model of choice for hippocampal studies, and particularly the rat hippocampus has yielded most of the detailed quantitative data we have about this structure. In 1990, Amaral and coworkers produced a comprehensive model of hippocampal neuronal populations and their connectivity in the rat based on the knowledge available at the time, a seminal work that became the foundation for the majority of models of the hippocampal network up to date (Treves and Rolls, 1994; Patton and McNaughton, 1995; Henze et al., 2002; Becker, 2005; Aimone et al., 2006, 2009; Leutgeb et al., 2007; Kempermann, 2011; Krueppel et al., 2011; Newman and Hasselmo, 2014; Chavlis et al., 2017; Berdugo-Vega et al., 2023; Borzello et al., 2023; Vandael and Jonas, 2024). Although some specific data have been updated from that original description (e.g. (Bannister and Larkman, 1995; Amaral and Lavenex, 2006; Bezaire and Soltesz, 2013), there has not been a comprehensive update including all the new information available regarding neuronal populations and the specifics of their connectivity. Therefore, inspired by that pioneer work, we have updated the model including significant developments described in recent years. In this manuscript we focus on the perforant projection to the dentate gyrus, updating the cytoarchitecture and connectivity of the entorhinal cortex (EC) and the dentate gyrus (DG), showing that their connectivity is much more divergent and richer than currently assumed.

The EC is the gateway for the cortical connections of the hippocampus, preprocessing and selecting salient and contextual information about the external world essential for the cognitive functions of the hippocampus (Kerr et al., 2007; Nilssen et al., 2019). In murine rodents, the EC has two major domains with differences in cytoarchitecture and functional profile: the lateral and medial entorhinal cortex subdivisions (LEC and MEC respectively), that correspond to areas 28a and 28b originally described by Brodmann (Brodmann, 2005, 2005; Amaral et al., 2007; Witter, 2007). The MEC conveys spatially specific information (Fyhn et al., 2004; Hafting et al., 2005), while the LEC seems to provide non-spatial contextual information related to odors, objects, temporal coordinates and emotional significance (Fyhn et al., 2004; Kerr et al., 2007; Tsao et al., 2018; Ohara et al., 2019).

The EC is a mesocortical area characterized by the absence of layer IV neurons that create a hypocellular layer (lamina dissecans) that separates upper layers II and II from deep layers V and VI. Entorhinal cortex projections to the hippocampus initiate in upper layers and include two main pathways: the perforant pathway that originates from layer II and projects mostly to the ipsilateral DG, CA3 and CA2, and the alvear or temporo-ammonic pathway that originates in layer III and terminates in CA1 and the subiculum bilaterally (Steward, 1976). In this manuscript we will focus on the perforant projections to the DG, leaving the rest of the perforant projections and the alvear pathway for future analyses.

The perforant pathway to the DG originates in EC layer II, terminates in the outer two thirds of the DG molecular layer (ML) and represents the vast majority of excitatory input into the dendrites of granule cells (GC) (Matthews et al., 1976; Amaral et al., 1990). In addition to EC layer II neurons, a small number of neurons from deep EC layers contribute to the perforant pathway but will not be considered here (for details see (Cappaert et al., 2015; Witter et al., 2017; Ohara et al., 2019). The perforant pathway is composed of two main components: the medial perforant pathway, originating from MEC and projecting to the medial molecular layer (MML) of the DG, and the lateral perforant pathway, originating from LEC and terminating mostly in the outer molecular layer (OML). This pathway has an additional level of topographic organization, as EC neurons close to the rhinal fissure connect more strongly to the septal hippocampus while ventral EC neurons have stronger connection with more temporal hippocampal levels (Canto et al., 2008; Kobro-Flatmoen and Witter, 2019), although for simplicity, we will not consider this gradient in the model.

Current models of the perforant pathway to the DG describe 200,000 EC layer II neurons projecting to 1 million GCs (Squire et al., 1989; Amaral et al., 1990; O’Reilly and McClelland, 1994; Patton and McNaughton, 1995; Becker, 2005; Aimone et al., 2006, 2009; Leutgeb et al., 2007; Newman and Hasselmo, 2014; Chavlis et al., 2017; Borzello et al., 2023), with a divergence of 1:5 that has been proposed to facilitate sparse encoding of information that facilitates pattern separation in the DG (O’Reilly and McClelland, 1994; Treves and Rolls, 1994). The current model also predicts that each EC neuron establish about 18,000 synapses with GCs and thus could contact up to ∼2% of the GC population, and in turn each GC receives on average about 4,000 synapses from EC neurons, meaning each GC may receive input from up to 2% of layer II EC neurons (Squire et al., 1989; Amaral et al., 1990; Patton and McNaughton, 1995; Derrick, 2007).

Our model incorporates a number of updates, starting with more precise estimates of the number EC layer II neurons obtained with design-based stereology, showing that males have approximately 116,000 neurons in layer II: about 70,000 in MEC and 46,000 in LEC (Arellano and Rakic, 2026). Another relevant update involves the revised definition of layer II in the LEC subfield based on molecular characterization of layer II neurons, that reshapes the traditional definition of Cajal and Lorente de No by dividing the layer into two sublayers: layer IIa, corresponding to the traditional definition of layer II, and layer IIb corresponding to the superficial aspect of what was considered traditionally as layer III (**Fig 1**). A third major update includes considering recent descriptions that the perforant pathway does not originate from all layer II neurons, but from a subpopulation of neurons expressing reelin that represents about half the neuronal population of layer II or about 80,000 neurons, ∼35,000 in MEC and ∼45,000 in LEC. Finally, although it is well known that the projections of MEC and LEC are segregated into the MML and OML of the dentate gyrus, this separation has not been considered when calculating the divergence of the connectivity between those entorhinal areas and the DG. When all those updates are combined, we obtain much higher divergence (1:20-1:30) in these projections, changing the calculus of the potential involvement of the DG in pattern separation.

**Figure 1.**
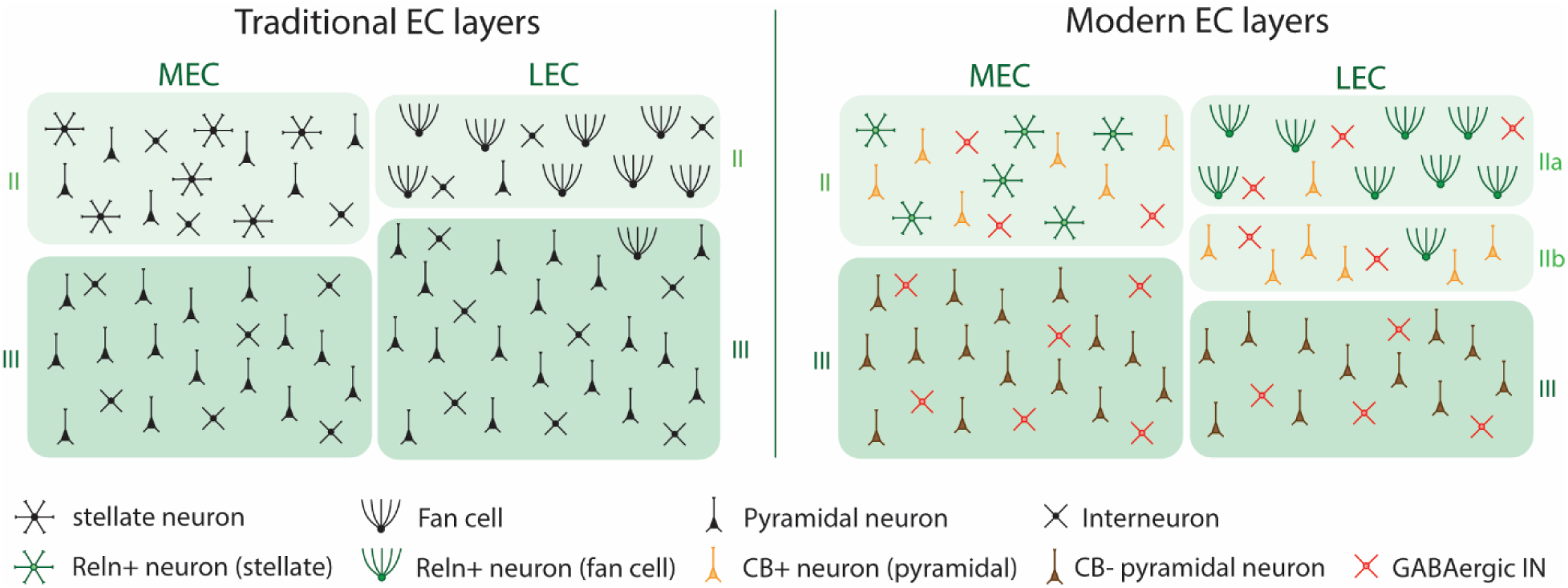
Changes in the cytoarchitectonic definition of superficial EC layers. The schematic illustrates the traditional definition of EC layer II from Cajal and Lorente de NO based on cytoarchitecture, that described a rim of relatively large cells, frequently described as stellate and fan cells with intermingled pyramidal neurons. The modern definition based on molecular characteristics describes two major populations of excitatory neurons: Reln+ and CB+, in addition to GABAergic inhibitory neurons. Reln+ neurons typically have stellate morphology in MEC and fan shape in LEC, while CB+ cells are typically pyramidal but see text for details on morphology. The modern characterization modifies the layering of LEC, with the traditional layer II populated mostly by Reln+ neurons becoming layer IIa, while the superficial aspect of traditional layer III, enriched in CB+ pyramidal neurons, becomes layer IIb.

The model also updates the number of excitatory inputs received by GCs, that we estimate to ∼11,000 per cell, with about 8,000 synapses originating from the EC. We also predict that each of the ∼80,000 layer II neurons from the EC projecting to the DG will establish about 80,000-115,000 synapses with GCs that represent about 75% of the excitatory input to GCs. We also model the redundancy in the connectivity, that according to available data might be about 25%, meaning that about 25% of the synapses in each GC originate from the same EC neuron, and therefore each EC neuron might contact 68,000-86,000 GCs. And reciprocally, each GC may receive input from 6,000 EC neurons (75% of 8,000 synapses).

In addition, we consider recent data obtained in mice indicating that septal mossy cells, in addition to their associational and commissural projections to the inner molecular layer, also project ipsi- and contralaterally to the MML, although it is not yet known if those projections target GCs or GABAergic interneurons (Botterill et al., 2021; Houser et al., 2021). Although those projections have not been described in rats, we estimate that ∼10,000 mossy cells might be responsible for that projection if present in rats. Since the target of these connections is not clear, we estimate that they might establish less than ∼200 synapses with each GC, that represents up to 5% of the synaptic input to GCS through the MML and suggests it will not significantly decrease MEC input to GCs. Conversely, if we consider that the main target of mossy cells in the MML are GABAergic interneurons, then we estimate that each interneuron might receive ∼1,600-6,400 excitatory synapse, which will represent a significant source of inhibition that could play a role in the regulation of GC activity.

Overall, the revised model of the perforant pathway to the DG presented here provides a number of updates regarding the cytoarchitecture, neuronal populations and connectivity ratios between EC layer II neurons and GCs that change our current understanding of the pathway, revealing much higher divergence than previously known, and much richer connectivity with GCs, with EC neurons establishing about 100,000 synapses with GCs, and connecting with 68,000-86,000 individual GCs that would represent 7-9% of the GC population, and each GC receiving input from ∼6,000 EC layer II neurons, that would represent ∼8% of the EC neurons projecting to the DG.

## Results

### Number of neurons in EC layer II

The overall entorhinal cortex population in the rat might be around 700,000 neurons (Arellano and Rakic, 2026). Upper layers II and III are the source of hippocampal projections, but for this manuscript, we are interested in layer II, the origin of the perforant path projecting to the DG, CA3 and CA2, although we will focus our analysis on the perforant pathway to the DG, leaving the perforant projections to CA3 and CA2 and the alvear pathway from layer III to CA1 and the subiculum for future studies.

Stereological studies using design-based stereology have shown that layer II contains approximately 116,000 neurons in males and 100,000 neurons in females (Arellano and Rakic, 2026). This sexual dimorphism in the population of layer II is expected, as it has been consistently described that the male hippocampus is larger, heavier and containing more neurons than the female hippocampus (reviewed in (Arellano and Rakic, 2026)). Since most hippocampal studies have favored the use of males to avoid the influence of cycling sexual hormones in the physiology of the hippocampal formation (Gould et al., 1990; Zhang et al., 2008), most available data on neuronal populations and connectivity comes from males, and for congruency, we focused our analysis on the number of neurons in layer II of males. However, there is no data for the separate population of MEC and LEC in males, and thus we estimated it from the total layer II population, using the ratio of 60:40 for MEC and LEC consistently reported in females (Arellano and Rakic, 2026). Thus, layer II in males might have about 70,000 neurons in MEC, and about 46,000 neurons in LEC.

A caveat of the above estimates is that they are based on the traditional definition of layer II by Cajal and Lorente de No, who described layer II as a narrow superficial rim mostly populated by large stellate-shaped neurons with some intermingled pyramidal cells (Kobro-Flatmoen and Witter, 2019). More recent molecular characterization of the neurons in EC layer II has shown it is composed of two main populations: reelin expressing (Reln+) and Calbindin expressing (CB+) neurons (Varga et al., 2010; Ray et al., 2014; Tang et al., 2014; Leitner et al., 2016; Ohara et al., 2019; Omholt et al., 2024), and while those two populations are intermingled in MEC layer II, they tend to segregate in LEC, where Reln+ cells occupy the narrow rim of relatively large cells originally identified as layer II that is now designated layer IIa, while the smaller CB+ pyramidal-like neurons are located subjacent to layer IIa, in a territory that was traditionally considered part of layer III and is now designated layer IIb (**Fig. 1**) (Kobro-Flatmoen and Witter, 2019; Ohara et al., 2019; Vandrey et al., 2020; Omholt et al., 2024). Thus, while there are no changes to the definition of MEC layer II and we can assume it contains 70,000 neurons as described above, the population of 46,000 neurons described in LEC would refer to the population of layer IIa. There are no available estimates for the number of neurons in layer IIb and the total population of (modern) LEC layer II, but we will obtain an estimate in the next section. Another consideration is that EC layer II contains 13-20% of GABAergic interneurons (Kumar and Buckmaster, 2006; Tang et al., 2014; Leitner et al., 2016), that need to be factored to obtain the population of excitatory neurons projecting to the DG.

### Number of EC layer II neurons projecting to the hippocampus

As indicated above, EC layer II excitatory neurons belong to two main populations: Reln+ and CB+ neurons (Varga et al., 2010; Ray et al., 2014; Tang et al., 2014; Leitner et al., 2016; Ohara et al., 2019; Omholt et al., 2024). However, only Reln+ neurons in MEC and LEC are the origin of the perforant pathway to the DG, CA2 and CA3 (Varga et al., 2010; Kitamura et al., 2014; Leitner et al., 2016; Vandrey et al., 2020) while CB+ have more segregated projections. MEC CB+ neurons send mostly intrinsic EC projections and a minor hippocampal projection through the alvear pathway targeting interneurons in the stratum lacunosum of CA1 that exert feed-forward inhibition over CA1 pyramidal neurons (Kitamura et al., 2014) while LEC CB+ neurons send a smaller projection to CA1 stratum moleculare, but mostly ipsi- and contralateral projections, specially to the EC but also to a variety of limbic areas such as olfactory structures, amygdala and ventro-medial prefrontal cortex (Varga et al., 2010; Kitamura et al., 2014; Fuchs et al., 2016; Leitner et al., 2016; Ohara et al., 2019).

Besides the modern classification by molecular features, layer II neurons in MEC have been traditionally classified based on morphology and physiology, primarily into stellate and pyramidal cells (Schwartz and Coleman, 1981; Alonso and Klink, 1993; Klink and Alonso, 1997; Gatome et al., 2010). Reelin expressing neurons are commonly identified as stellate cells and CB+ neurons as pyramidal cells (Kitamura et al., 2014; Ray et al., 2014; Tang et al., 2014; Sürmeli et al., 2015). However, such morphological segregation is not clear cut (Canto and Witter, 2012; Fuchs et al., 2016; Witter et al., 2017; Ohara et al., 2019; Omholt et al., 2024), as for example, RE+ cells in LEC typically show a fan-like morphology and include a certain proportion of pyramidal and intermediate-shaped neurons (Nilssen et al., 2019; Vandrey et al., 2020; Omholt et al., 2024). Thus, it seems that Reelin expression, rather than morphology, may be the best criterion to calculate the number of EC neurons projecting to the hippocampus.

Reelin expression is characteristic of EC layer II across mammalian species, with a consistent gradient of expression from lateral (rhinal sulcus) to medial (Drakew et al., 1998; Pesold et al., 1998; Pérez-García et al., 2001; Martínez-Cerdeño et al., 2003; Ramos-Moreno et al., 2006; Kobro-Flatmoen and Witter, 2019). Quantitative studies in rat have focused on MEC, providing the proportions of Reln+ and CB+ neurons in layer II (Varga et al., 2010; Tang et al., 2014), showing that ∼50% of neurons are Reln+ and 36% are CB+ after adjusting for a ∼14% GABAergic population (Kumar and Buckmaster, 2006; Tang et al., 2014)(**Table 1**). Thus, Reln+ neurons seem to represent the larger population, in agreement with data showing that most neurons in MEC layer II project to the DG (Kitamura et al., 2014). Conversely, (Ray et al., 2014) described a majority of CB+ neurons (60%) in the cell clusters of MEC layer II. However, that quantification excluded neurons interspersed between the clusters that are typically Reln+ (Kitamura et al., 2014) and may not be representative of the overall layer. Thus, we can estimate that Reln+ neurons represent ∼35,000 neurons (49% of 70,000) in MEC layer II (**Table 2**).

**Table 1:** Estimates of the proportion of reelin (Reln), calbindin (CB) and GABAergic interneurons in the MEC of rats. *Note: (Varga et al., 2010) original proportions of 53% Reln+ and 44% CB+ did not consider GABAergic interneurons. Here we show adjusted percentages for 14% interneurons (see text).

| Reference | Reln (%) | CB (%) | Reln/CB (%) | GABA (%) |
| --- | --- | --- | --- | --- |
| Varga et al., 2010* | 45 | 38 | 2.8 | 14 |
| Tang et al., 2014 | 53 | 34 | - | 13 |
| Average MEC | 49 | 36 | - | 14 |

**Table 2.** Neuronal populations in EC layer II. Estimates of Reln+ and CB+ neurons in areas MEC and LEC of males and females. Male data are estimated as 15% more neurons than females. Reln+ and CB+ neurons in MEC calculated with data from table 1. LEC layer IIb CB+ neurons estimated assuming similar proportion than in MEC (∼36%). Total LEC IIb population and number of GABAergic interneurons in layer IIa and IIb estimated using data from mouse showing that CB+ neurons represent 70% of LEC layer II and GABAergic interneurons represent 13% in layer IIa and ∼20% in layer IIb (Leitner et al., 2016). Data is rounded for simplicity and totals might not match.

| Sex | Area | Neurons | Reln+ neurons | CB+ neurons | % GABA+ neurons | GABA+ neurons |
| --- | --- | --- | --- | --- | --- | --- |
| female | MEC | 60,000 | 30,000 | 22,000 | 14 | 8,400 |
|  | LEC IIa | 40,000 | 35,000 | 600 | 13 | 5,200 |
|  | LEC IIb | 37,000 | 4,000 | 26,000 | 20 | 7,300 |
|  | LEC | 77,000 | 39,000 | 27,000 | ~16 | 12,500 |
|  | EC | 137,000 | 69,000 | 49,000 |  | 20,900 |
| male | MEC | 70,000 | 35,000 | 25,000 | 14 | 9,800 |
|  | LEC IIa | 46,000 | 40,000 | 700 | 13 | 6,000 |
|  | LEC IIb | 42,000 | 5,000 | 30,000 | 20 | 8,400 |
|  | LEC | 88,000 | 45,000 | 30,000 | ~16 | 14,400 |
|  | EC | 158,000 | 80,000 | 55,000 |  | 24,200 |

Regarding LEC, it has been described that Reln+ neurons represent the vast majority of layer IIa (Leitner et al., 2016; Kobro-Flatmoen and Witter, 2019; Vandrey et al., 2020; Omholt et al., 2024) in agreement with immunohistochemical studies showing that virtually all neurons in the traditional LEC layer II (corresponding to layer IIa) express reelin (Martínez-Cerdeño et al., 2003; Ramos-Moreno et al., 2006).

Thus, we can assume that the ∼46,000 neurons estimated for traditional layer II in males, and layer IIa by the modern definition, are mostly Reln+. In mice, (Leitner et al., 2016) showed that 13% of layer IIa are GABAergic, meaning about 6,000 neurons will be GABAergic interneurons in rat, leaving about 40,000 excitatory Reln+ neurons and a small population of excitatory CB+ neurons (**Table 2**). Regarding layer IIb, we can speculate about its population, assuming it has the same proportion of CB+ neurons (36%) than MEC layer II, that would amount to 30,000 excitatory CB+ neurons. Other populations can be deducted using mice data from (Leitner et al., 2016): they describe that CB+ neurons represent 70% of the total population of layer IIb, implying layer IIb would have about 42,000 neurons in total, 8,400 of which (20%) would be GABAergic interneurons and 5,000 (12%) will be excitatory Reln+ neurons (Leitner et al., 2016)**(Table 2)**. It seems plausible that those 5,000 excitatory Reln+ neurons in layer IIb represent “ectopic” layer IIa neurons that also project to the hippocampus. Thus, combining them, males might have about 45,000 Reln+ neurons in LEC layer II projecting to the hippocampus.

In summary, the number of EC layer II neurons projecting to the DG in male rats can be estimated to be ∼35,000 neurons in MEC, and ∼45,000 neurons in LEC. (Kobro-Flatmoen et al., 2016) using transgenic male and female rats reported ∼121,000 Reln+ neurons in MEC layer II and 47,000 in LEC layer II (average of 3 and 6-month-old animals). While their estimate for LEC is about the same as ours, their estimate for MEC is almost 4-fold. Even after removing 6.6% potential Reln+ GABAergic interneurons (Leitner et al., 2016) there would be ∼113,000 excitatory Reln+ neurons in MEC that is still more than 3 times our estimate, and almost twice the overall stereological estimate for MEC layer II that includes CB+ neurons (Arellano and Rakic, 2026). Although some divergences are expected due to differences in the segregation of fields or because they used transgenic rats, we do not have an explanation for the marked difference in the MEC estimates and we did not use those data.

### Number of GC neurons

The dentate gyrus comprises two cellular layers: the granular cell layer (GCL), densely populated by GC somata whose dendrites ramify in the ML, and the polymorphic layer or hilus, populated by both excitatory hilar mossy cells and GABAergic interneurons. The GCL contains about 1 million neurons in males, mostly GCs but also GABAergic interneurons that however represent less than 1% (Czéh et al., 2013) and therefore for simplicity we will assume the GCL contains 1 million GCs. Also, modified GC-like neurons in the IML (semilunar cells of Cajal (Ramon y Cajal, 1893; Williams et al., 2007; Larimer and Strowbridge, 2010) and ectopic GCs in the hilus and CA3 (Amaral and Woodward, 1977; Martí-Subirana et al., 1986; Tóth and Freund, 1992; Szabadics et al., 2010) have been described, but they represent small populations and therefore will not be further discussed here.

### EC input to GCs

Granule cell dendrites ramify in a fan-like fashion in the molecular layer (ML), a compartment that can be segregated in three main strata according to the input it receives. The inner third or inner molecular layer (IML) receives mostly associational and commissural input originating from ipsi- and contralateral hilar mossy cells and from subcortical structures, with a minor contribution from other fields such as CA3 and subiculum (Cappaert et al., 2015). The outer two thirds of the ML receive almost exclusively excitatory synapses from the entorhinal cortex (Matthews et al., 1976; Amaral et al., 1990) segregated in two laminae: the medial molecular layer (MML) that receives mostly input from the MEC and the outer molecular layer (OML) that receives mostly input from the LEC. These entorhinal projections have a small contralateral contribution (Goldowitz et al., 1975; Zimmer and Hjorth-Simonsen, 1975; Steward, 1976), that we will not consider for the present model. Additionally, recent reports have shown that the MML might receive also a sizable quantity of excitatory synapses from septal mossy cells (Botterill et al., 2021; Houser et al., 2021), although the target of those synapses has not been determined.

Stereological estimates have described 10,000-11,000 presumed excitatory synapses per GC (West and Andersen, 1980; Cardoso et al., 2008a; Thind et al., 2010), mostly in spines (98%) and a small fraction (2%) in dendritic shafts (Matthews et al., 1976; Thind et al., 2010). The IML was described to receive about 25% of the input or ∼2,700 synapses, while the outer two thirds corresponding to MML and OML would receive 75% or about 8,200 (Crain et al., 1973; Geinisman et al., 1991, 1992; Cardoso et al., 2008b; Thind et al., 2010) (**table 3**). Also, (Andrade et al., 2002) described about 820 axo-spinous synapses per GC in the MML and OML, suggesting it could be a typo and might correspond to 8,200 axo-spinous synapses per GC in the MML and OML, a figure that will agree with the other estimates. Finally, another study described much larger number of synapses (∼55 billion; (Popov and Stewart, 2009)) implying each granule cell would receive about 55,000 excitatory synapses, or 17 spines/micron (50,000 spines/3,200 um; see below), a spine density that is obviously too high and therefore will not be considered here.

**Table 3.**
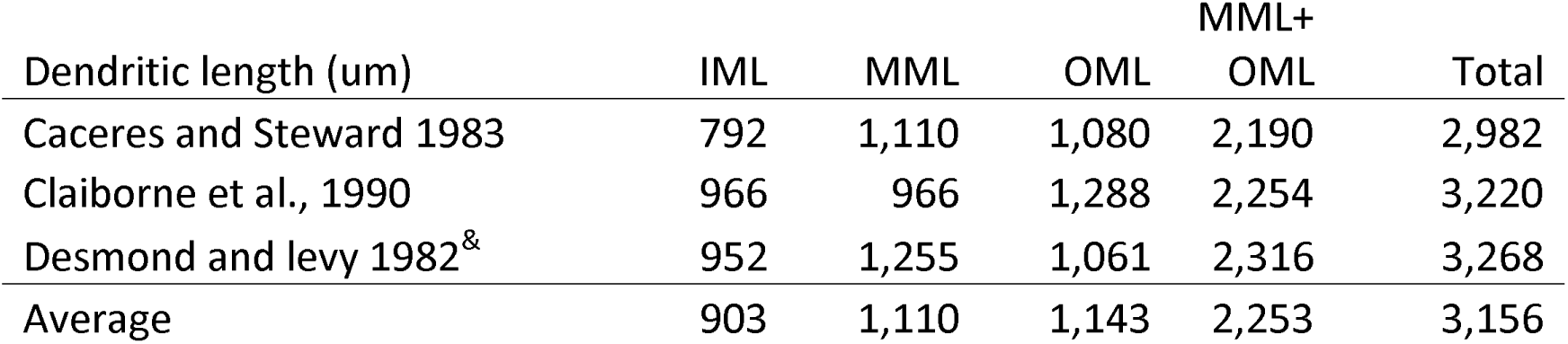
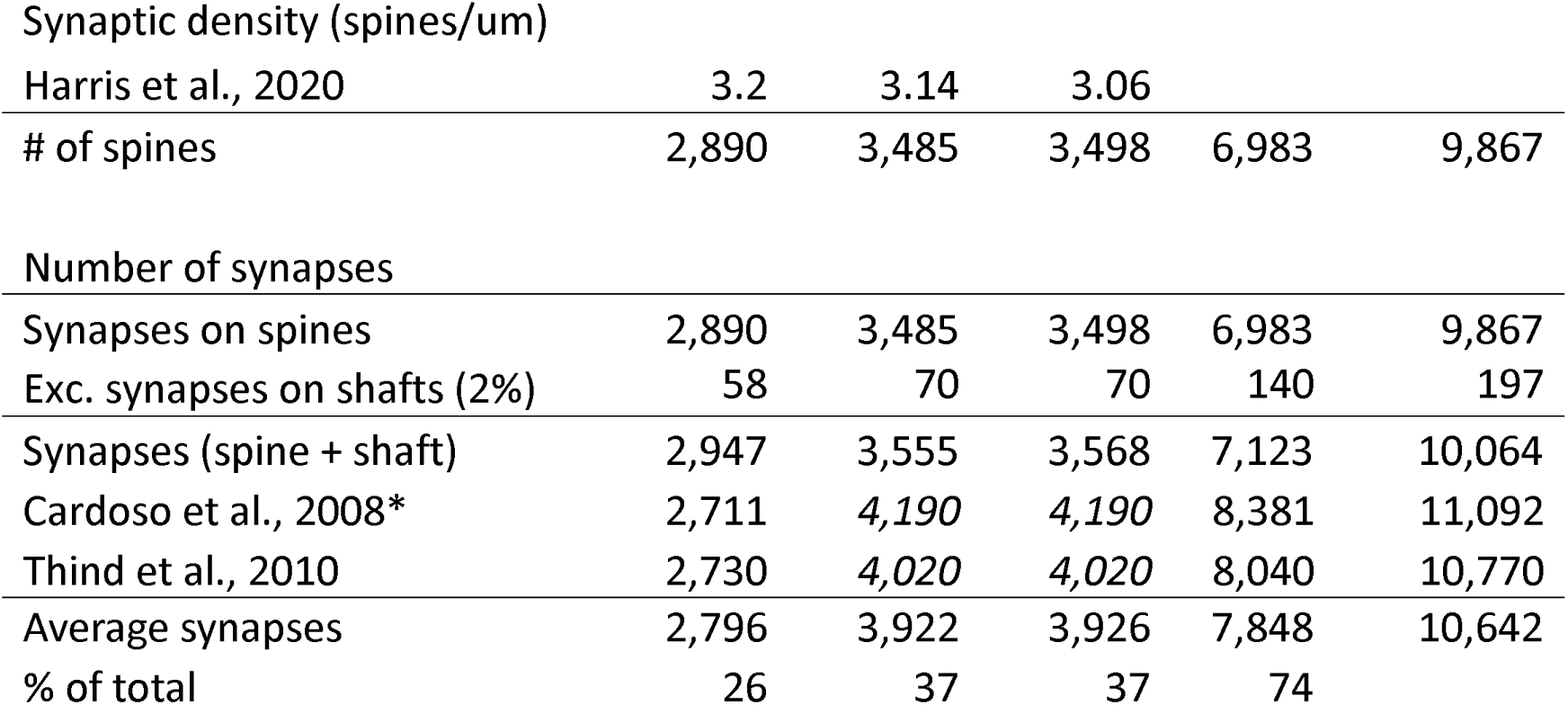
Estimates of number of synapses in the strata of the ML obtained from spine estimates combining dendritic length and synapse density and stereological studies. Both approaches render quite similar estimates. All studies used male animals except Claiborne et al., 1990 that used both males and females. &Desmond and Levy 1982 reported a total dendritic length of 3,662 um for GCs, including 394 um in the GCL. Here we consider only the dendritic length in the ML. * Data for synapses on spines only. (Cardoso et al., 2008a) reported very few symmetric synapses, contrary to most available evidence, and thus axo-dendritic synapses, which would correspond mostly to symmetric synapses are not included here.

Another approach to estimate the input of GCs is through the number of dendritic spines, that receive most of the excitatory input (Matthews et al., 1976; Thind et al., 2010). Detailed studies in rat have shown that the dendritic length of GCs ranges between 2,500-4,600 um, with an approximate average of ∼3,200 um (Desmond and Levy, 1982; Caceres and Steward, 1983; Claiborne et al., 1990; Cannon et al., 1999; Pierce et al., 2011; Buckmaster, 2012; Cole et al., 2020) (**Table 3**). Longer dendritic trees in GCs from the suprapyramidal blade compared to the infrapyramidal blade have been reported (Rahimi and Claiborne, 2007), but for simplicity, we will consider only the average here. Total dendritic length is distributed quite homogeneously in the three strata, with approximately ∼1000 microns in each compartment (**Table 3**). Regarding the density of spines, most studies using brightfield microscopy describe values between 1-2 spines per micron e.g.: (Desmond and Levy, 1985; Gould et al., 1990; Cole et al., 2020; Puhahn-Schmeiser et al., 2022; Smilovic et al., 2022). Although quantification of spines at the optical microscope level is useful to establish comparisons between species or experimental conditions, it is not an accurate method to obtain the total number of spines, as it is prone to underestimation due to resolution limitations and the visual obstruction created by the dendrite and other spines (Feldman and Peters, 1979). In this regard, electron microscope (EM) three-dimensional analysis produces the most accurate counts, although with the limitation of small sampling size. High voltage EM analysis showed densities between 2 and 3.4 spines/micron (Hama et al., 1989), and detailed dendrite reconstruction, the most accurate method, showed similar but slightly higher spine densities that were similar in the three strata: 3.2 spines/micron in the IML, 3.14 in the MML and 3.06 in the OML (Harris et al., 2022). Thus, combining dendritic length and EM based spine density, it can be estimated that on average, each GC has ∼9,900 spines, each typically receiving 1 excitatory synapse, meaning GCs receive ∼9,900 axo-spinous synapses (**Table 3**), and another ∼200 (2%) on dendritic shafts (Matthews et al., 1976; Thind et al., 2010; Bromer et al., 2018) to a total of 10,100 excitatory synapses, very similar values to those described above from stereological quantification studies (**Table 3**).

All quantitative hippocampal models we encountered described entorhinal projections to the DG generically, originating from EC neurons, without differentiating between LEC and MEC populations. However, these divisions, corresponding to the original subfields 28a and 28b of Brodmann (Brodmann, 2005; Witter, 2007) exhibit different cytoarchitecture, each have segregated connections with the entire DG (Hjorth-Simonsen, 1972; Hjorth-Simonsen and Jeune, 1972; Witter, 2007) and are involved in different information modalities, with MEC processing mostly spatial information, and LEC dealing with contextual information (Fyhn et al., 2004; Hafting et al., 2005; Kerr et al., 2007; Tsao et al., 2018; Ohara et al., 2019). Thus, we considered them as separate EC fields to estimate their connectivity with the DG, with MEC exhibiting 70,000 neurons and LEC 46,000 neurons in layer II of male rats (and 60,000 and 40,000 in females).

Spine analysis also provides separate data for IML, MML and OML input, indicating that MEC and LEC projections to GCs are very similar, each producing about 3,500 excitatory synapses or 35% of the total excitatory input to GCs, while the IML receives about 2,950 excitatory synapses in the IML or about 30% of the total. Those percentages are similar to the 25% for the IML and 75% for the combined MML and OML revealed by stereological estimates of synapses. Averaging spine estimates and stereological estimates suggest a total of ∼10,640 synapses per GC, with ∼2,800 synapses (26%) in the IML; and ∼3,920 synapses (37%) in both the MML and OML (**Table 3**) that we will round to 4,000 for the MML and OML for simplicity (**Fig. 2**).

**Figure 2.**
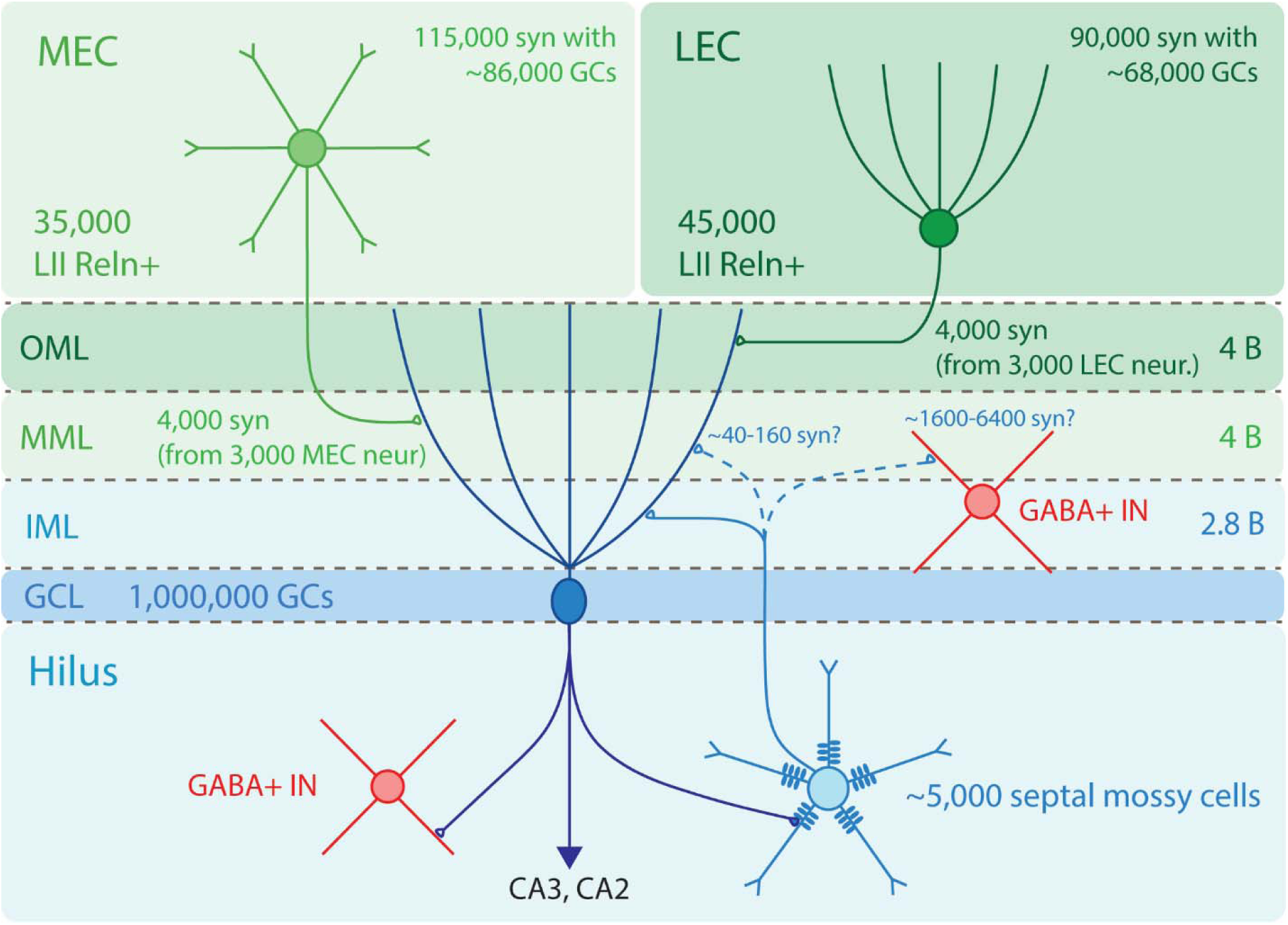
Schematic representation of the connectivity of GCs. About 35,000 MEC neurons send axons to the MML establishing about 115,000 synapses with GCs but contacting about 86,000 GCs due to 25% redundance in the connectivity. Also, 45,000 LEC neurons establish about 90,000 synapses with GC dendrites in the OML, contacting about 68,000 GCs. In turn, each GC is contacted by about 8,000 EC synapses, ∼4,000 synapses in MML from about 3,000 MEC neurons, and ∼4,000 synapses in OML from about 3,000 LEC neurons. In addition, each GC receives about 2,800 synapses in the IML from ipsi- and contralateral mossy cells. And there is a possibility that GCs receive up to ∼160 synapses from septal mossy cells in the MML (or those synapses may contact Inhibitory interneurons, establishing up to 6,400 synapses with each interneuron). Abbr: B: billion; GABA+ IN: GABAergic interneuron; GC: granule cells; GCL: granular cell layer; IML: inner molecular layer; LEC: lateral entorhinal cortex; LII Reln+: layer II neurons expressing reelin; MEC: medial entorhinal cortex; MML: medial molecular layer; neur: neuron; OML: outer molecular layer; syn: synapse.

Additionally, occasional dendrites from some mossy cells (∼12%) enter the molecular layer, typically to the IML, but in rare cases reaching MML and OML (Amaral, 1978; Scharfman, 1991; Buckmaster et al., 1992, 1993; Scharfman and Myers, 2012) and mossy cells have been recorded to respond directly to EC stimulation (Scharfman, 1991; Scharfman and Myers, 2012). Quantitative data on this connectivity is lacking, although it seems that it might represent a small ratio of mossy cell input and is unlikely to significantly alter the estimates of EC connectivity with GCs provided above, and thus we will not consider it further in the analysis.

### GC input from EC (MML and OML)

As described in the EC section, about 35,000 Reln+ neurons in MEC layer II and 45,000 in LEC layer II project to 1 million GCs in the dentate gyrus, with divergence ratios around 1:30 and 1:20 for MEC and LEC, respectively (**Fig. 2**). In addition to the projections from layer II, a small number of neurons from deep EC layers contribute to the perforant pathway but they will not be considered here. For further information see (Cappaert et al., 2015; Witter et al., 2017, 2017; Ohara et al., 2019).

Projections from MEC terminate in the MML and those from LEC terminate in the OML. EC input is considered virtually the only input arriving to those strata, although in mice it has been described an undetermined number of excitatory synapses in the MML that seem to originate from hilar cells located in the septal hippocampus (Botterill et al., 2021; Houser et al., 2021). Each GC neuron receives on average about ∼8,000 entorhinal connections, divided evenly between the MML (∼4,000 synapses) and OML (4,000 synapses) (**Table 3, 4; Fig. 2**). Thus, GCs receive about 4 billion synapses (3.92 billion) in the MML from MEC layer II Reln+ neurons, and another 4 billion in the OML from LEC layer II Reln+ neurons. This also implies that each of the 35,000 MEC layer II Reln+ neurons will establish on average ∼115,000 synapses with GCs, while each of the 45,000 from LEC would establish ∼90,000 synapses with GCs **(Table 4**, **Fig. 2)**.

**Table 4:** Quantitative synaptic connectivity between the EC and GCs. The number of targeted GCs per EC neuron and the number of EC neurons synapsing on each GC are calculated as 75% of the total descried, assuming a redundancy of 25%. Figures have been rounded for simplicity. For details, see text. EC: entorhinal cortex; GC: granule cells; LEC: lateral entorhinal cortex; LII: layer II; MEC: medial entorhinal cortex; MML: medial molecular layer; ML: molecular layer; OML: outer molecular layer; syn: synapse # About 12% of mossy cells have 1-2 dendrites in the MML of the DG, suggesting larger overall number of EC synapses in MML. Also, septal mossy cells send projections to the MML in mice that might contact GCs, although it might represent a small portion of the GC input, not changing significantly the rate of EC input to GCs (see text).

| EC field | # EC neurons<br>proj to<br>DG | DG<br>Target | Popul<br>ation<br>diver<br>gence | EC syn<br>in ML | GCL<br>syn/ EC<br>neuron | Targeted<br>GCs/EC<br>neuron (%) | Syn/<br>GC | EC neurons<br>synapsing<br>on each GC<br>(%) |
| --- | --- | --- | --- | --- | --- | --- | --- | --- |
| MEC LII | 35,000 | MML<sup>#</sup> | 1:30 | 4 billion | 115,000 | 86,000 (8.6) | 4,000 | 3,000 (8.6) |
| LEC LII | 45,000 | OML | 1:20 | 4 billion | 90,000 | 68,000 (6.8) | 4,000 | 3,000 (6.6) |

The above calculations are based on data from males. Since, as discussed above, the evidence suggests that both EC and GC populations might be ∼15% smaller in females, the overall neuronal and connectivity ratios between EC layer II neurons and GCs might be similar in both sexes despite their differences in size and neuronal populations.

### Redundancy in the connectivity

Given the large EC input to GCs, it is relevant to consider how redundant it is, or in other words, how many GCs are contacted by a single EC axon. This is an important concept already emphasized by (Amaral et al., 1990), as the connectivity might be governed by maximum spread (divergence), each EC neuron connecting only once with each GC, so each MEC neuron might contact 115,000 GCs and each LEC neuron about 90,000 GCs (12 and 9% of the GC population), or, conversely, connectivity could be more selective, with EC neurons establishing multiple connections with less GCs, for example an average of 10 synapses per GC, and then each MEC neuron will contact on average 12,000 GCs and each LEC neuron about 9,000 GCs (1.2 and 0.9% of the GC population).

Detailed EM analysis of discrete GC dendritic segments have shown examples of axons contacting 3 spines from the same dendrite (Popov and Stewart, 2009) and more generally, that about 13% of synapses on GC spines are established by an axon that contacted another spine in the same dendrite (26 spines (13 pairs) out of 96; (Bromer et al., 2018)). If we assume that similar redundancy (∼13%) is present between different dendritic segments of the same cell, then about 25% of the synapses in each GC would be redundant, and therefore each MEC neuron will contact ∼86,000 GCs (75% of 115,000 GCs) and each LEC neurons about 68,000 GCs (75% of 90,000 GCs), and conversely each GC would receive input from ∼3000 MEC neurons (75% of 4000) and ∼3,000 LEC neurons (75% of 4,000) (**Fig. 2**), that represent about 9% of the population of Reln+ excitatory cells in MEC and about 7% in LEC. Finally, another possibility to consider is that dendritic spines might receive more than one synapse, increasing the connectivity of each granule cell. However, available data indicates that only about 0.5% of spines receive more than one excitatory synapse (Popov and Stewart, 2009), a low percentage that would not change substantially the calculations obtained above.

In addition to excitatory cells, cells with GABAergic interneuron morphology in the EC have also been reported to send projections to the ML (Germroth et al., 1989b, 1989a; Fifkova et al., 1992) but they seem to target mostly GABAergic interneurons (Melzer et al., 2012). Also, hippocampal interneurons project to the EC (Melzer et al., 2012).

### Hilar mossy cell projections to the DG

The hilus corresponds to the polymorphic layer of the dentate gyrus and contains two major types of neurons: excitatory mossy cells and inhibitory GABAergic interneurons. Stereological studies describing the number of neurons in the hilus have usually quantified both populations together, as they are impossible to differentiate in cellular staining. Those studies have described about 50,000 hilar neurons (Arellano and Rakic, 2026), about 65% of which, or 33,000 are mossy cells (Buckmaster and Jongen-Rêlo, 1999). However, the distribution of mossy cells is quite different at septal or temporal levels. First, the space demarcated by the GCL is much larger temporally than septally, and therefore hilar cells are more numerous in the temporal half (68-73%) than in the septal half (27-32%) (Gaarskjaer, 1978; Rapp and Amaral, 1988; Buckmaster and Dudek, 1997; Buckmaster and Jongen-Rêlo, 1999; Patel and Bulloch, 2003; Jiao and Nadler, 2007). And second, the proportion of mossy cells from the total hilar cells is higher in the temporal half (65%) than in the septal half (45%), meaning that there might be about 7 times more mossy cells in the temporal half (86%) than in the septal half (14%) of the hilus (Patel and Bulloch, 2003; Jiao and Nadler, 2007).

Mossy cells in the hilus send most of their projections to the IML of the DG both ipsi and contralaterally, representing the second largest source of excitatory input to GCs after the perforant pathway. But in addition to that major projection to the IML, early descriptions of the mossy cell axons described a low fraction of axonal length (∼1-4%) distributed also in the MML, and scarcely in the OML (Buckmaster et al., 1992, 1996). More recent studies using molecular characterization of mossy cells in mice have shown projections to the MML across the septotemporal axis that originate from mossy cells located in the septal hippocampus, while mossy cells located at mid septotemporal and temporal levels show the classical innervation exclusively in the IML (Botterill et al., 2021; Houser et al., 2021). It is not known if those projections target preferentially GCs or GABAergic interneurons, and there is no quantitative data regarding how many synapses they establish in the MML. However, if they terminate preferentially on GCs, these excitatory projections are expected to terminate in dendritic spines of GCs, competing with perforant input. Therefore, we will calculate a rough estimate of their potential connectivity with GCs to figure out if they could alter the estimated input from MEC to the MML described above, and we will also estimate their potential connectivity with GABAergic interneurons.

### Estimate of hilar mossy cell projections to the MML

The projections of mossy cells to the molecular layer are restricted to the septal half of the hippocampus, so we can speculate it may involve up to ∼15% of mossy cells or 5,000 neurons (see above). Since the projection seems to be also commissural (Botterill et al., 2021; Houser et al., 2021), we could estimate it involves about 10,000 septal mossy cells from both hemispheres. There is no data about their potential targets or their synaptic coverage, so they can synapse on dendrites of GCs, on GABAergic interneurons, or in a combination of both. Nonetheless, we can speculate, from the dense axonal plexuses reported, that it might represent about 10-20% of the 4 billion excitatory synapses in the MML, or 400-800 million synapses overall. However, that would imply that each of the 10,000 septal mossy cells should establish on average 40,000-80,000 synapses in the MML, numbers that are significantly higher than their average synaptic output to the IML. Thus, it seems more likely that they might establish a much smaller proportion of synapses in the MML, in the range of 1-4% described in previous studies (Buckmaster et al., 1992, 1996), that means a total of 40-160 million synapses or ∼4,000-16,000 synapses per mossy cell, that still represents up to half their number of synapses in the IML. If those synapses target preferentially GCs, then each GC will receive on average 40-160 synapses (**Fig. 2**), a very low proportion of the total. This low contribution of non-EC input is supported by data showing that complete lesion of the EC produces degeneration of 86% of synapses in both OML and MML, leaving symmetric, inhibitory synapses unaltered (Matthews et al., 1976). Considering inhibitory synapses may represent 6-22% in the ML (Gottlieb and Cowan, 1972; Crain et al., 1973; Halasy and Somogyi, 1993; Thind et al., 2010), it seems there might be little additional input besides that from the EC.

Another possibility is that those projections target preferentially GABAergic interneurons. There is no data available on the number of interneurons with dendrites in the MML, but we can speculate that this number could be around 25,000 interneurons: 20,000 interneurons with soma in the GCL or MML and about one third (∼5,000) of the ∼17,000 hilar interneurons (Buckmaster and Jongen-Rêlo, 1999; Shi et al., 2004; Thind et al., 2010; Czéh et al., 2013). And if we assume a similar number of mossy cells synapses (40-160 million) as calculated for GCs, that would represent ∼1,600-6,400 excitatory synapses per GABAergic interneuron (**Fig. 2**), that fits in the range of ∼1,600-15,000 excitatory synapses estimated for GABAergic interneurons in other hippocampal fields such as CA1 (Gulyás et al., 1999).

## Discussion

The aim of the present work is to review and update the model of connectivity of excitatory circuits in the hippocampal formation. The seminal models by (Amaral et al., 1990) or (Patton and McNaughton, 1995) produced useful blueprints of hippocampal connectivity that were later updated in some of their parameters (Bannister and Larkman, 1995; Amaral and Lavenex, 2006; Dyhrfjeld-Johnsen et al., 2007; Morgan et al., 2007; Bezaire and Soltesz, 2013), but a full review incorporating all new available data on hippocampal connectivity is missing, and we aimed to fill that void. To this end, we used updated populations of hippocampal neurons (Arellano and Rakic, 2026) and detailed information on their connectivity that has become available in the last three decades to produce an updated model of connectivity for the hippocampal formation of the male rat. For conciseness, in this manuscript we focus on the perforant pathway to the DG, leaving the rest of the EC projections to the hippocampus, and the remaining hippocampal connectivity for future analyses.

As mentioned, the model is based on data from male rats. The specification of species and sex is important, because although the basic structure and connectivity of the hippocampal formation is conserved across mammals (with the exception of cetaceans, that show marker hippocampal regression), there are reported differences in connectivity and extent of projections across species, e.g.: (van Groen et al., 2002, 2003; Witter, 2007). Also, sex differences, at least in the number of neurons in hippocampal fields, have been reported (Arellano and Rakic, 2026), and therefore caution should be applied before generalizing the present model to female rats and to other species, even close ones as the mouse. In this regard, it can be argued that it might be more useful to model the mouse hippocampus, as mice have become the preferred animal model for research. We agree with that premise, but even when detailed data on mouse hippocampal circuits is increasing at fast pace, there is still much more detailed data published on synaptic connectivity of the rat hippocampus, allowing a more detailed and complete analysis. Nonetheless, the present model can be useful as a blueprint for comparative studies of connectivity between species, and ultimately with the human hippocampus.

We focused on the perforant pathway because its original description has been profoundly changed by a number of findings including updated populations of EC layer II neurons from stereological studies; revision of the cytoarchitecture of LEC layer II based on molecular features; redefinition of the neurons projecting to the hippocampus now restricted to neurons expressing Reelin; quantitative characterization of the population of GABAergic interneurons and finally segregation of fields MEC and LEC as independent sources of input. After considering all those factors, we describe a population of 35,000 neurons from MEC and 45,000 neurons from LEC projecting to the DG, substantially smaller populations than the traditional 200,000 neurons used in most models (Amaral et al., 1990; O’Reilly and McClelland, 1994; Patton and McNaughton, 1995; Becker, 2005; Aimone et al., 2006, 2009; Leutgeb et al., 2007; Newman and Hasselmo, 2014; Chavlis et al., 2017; Borzello et al., 2023), and changing significantly the divergence and synaptic relations of this projection, that is the source of cortical information to the DG.

The perforant pathway targets mostly GC dendrites in the ML of the DG. We estimated a population of 1 million GCs (Arellano and Rakic, 2026), matching the traditional figure used in most previous models e.g.: (Amaral et al., 1990; O’Reilly and McClelland, 1994; Treves and Rolls, 1994; Patton and McNaughton, 1995; Kempermann, 2011; Chavlis et al., 2017). We estimated the synaptic input to those GCs by calculating their number of dendritic spines using dendritic length and updated spine densities based on EM data, combined with data from stereological studies quantifying excitatory synapses in the ML. Both methods converge to show that GCs receive about 8,000 synapses from the perforant pathway, with MEC and LEC contributing evenly with about 4,000 synapses each. Then, the 1 million GCs receive about 4 billion synapses from MEC and another 4 billion from LEC, implying that each of the 45,000 MEC neurons establish about 96,000 synapses with GCs, and each of the 35,000 LEC neurons form about 115,000 synapses with GCs. We also estimated a redundancy in the connectivity of about 25%, meaning each GC would receive input from about 3,000 EC neurons from MEC and 3,000 from LEC, that represent 7-9% of the population of MEC and LEC (respectively) projecting to the DG. And reciprocally, each neuron from MEC will connect with 68,000 GCs, while each LEC neuron will connect with 86,000 GCs.

Overall, the current model presents an updated picture of the perforant pathway to the DG, showing that relatively small populations from two functionally and cytoarchitectonically distinct EC areas send very rich and extensive projections to the DG, with overall high divergence and extensive coverage of the GC population. The details and implications of these findings are analyzed in the next sections.

### EC Layer II neurons projecting to the hippocampus are a surprisingly small population that might rely on signal and frequency encoding

The EC receives cortical input from associative, limbic, and unimodal and multimodal sensory areas, and funnels that cortical information into the hippocampus, preprocessing and selecting salient and contextual information about the external world essential for the cognitive functions of the hippocampus (Kerr et al., 2007; Nilssen et al., 2019). Our analysis reports an updated population of 116,000 neurons in EC layer II in males and 100,000 neurons in females, that represent about half the 200,000 EC neurons originally described and reported in most models of hippocampal function. This difference can be explained considering that 200,000 was a rough estimate obtained from the product of the areal density of neurons in EC layer II and the surface area of the EC, originally reported with a question mark to emphasize it was an approximation (see fig 1 in (Amaral et al., 1990)).

All quantitative hippocampal models we encountered described entorhinal projections to the DG generically, originating from EC neurons, without differentiating between LEC and MEC populations. However, these divisions, corresponding to the original subfields 28a and 28b of Brodmann exhibit different cytoarchitecture (Brodmann, 2005; Witter, 2007), are involved in different information modalities, with MEC processing mostly spatial specific information (Fyhn et al., 2004; Hafting et al., 2005) and LEC dealing with non-spatial contextual information related to odors, objects, temporal coordinates and emotional significance (Fyhn et al., 2004; Kerr et al., 2007; Tsao et al., 2018; Ohara et al., 2019), and each have segregated connections with the entire DG (Hjorth-Simonsen, 1972; Hjorth-Simonsen and Jeune, 1972; Witter, 2007). Thus, functionally and in terms of connectivity, they can be considered separate EC areas with MEC exhibiting 70,000 neurons and LEC 88,000 neurons in layer II of male rats, the later after considering the redefinition of LEC layer II based on molecular features that now distinguishes layer IIa and IIb (Kobro-Flatmoen and Witter, 2019; Ohara et al., 2019; Vandrey et al., 2020; Omholt et al., 2024). Those populations would be 60,000 and 77,000 in females, respectively.

Another important update considered in the model is the molecular segregation of the two major populations of excitatory neurons in EC layer II into Reln+ and CB+ neurons (Varga et al., 2010; Ray et al., 2014; Tang et al., 2014; Leitner et al., 2016; Ohara et al., 2019; Omholt et al., 2024). Considering only Reln+ neurons project to the DG (Varga et al., 2010; Kitamura et al., 2014; Leitner et al., 2016; Vandrey et al., 2020), we obtained estimates of 45,000 and 35,000 Reln+ neurons in MEC and LEC, respectively, projecting to the DG.

As mentioned above, the EC receives rich and complex cortical information from many cortical areas, preprocessing and selecting salient and contextual information about the external world that is directed to the hippocampus to create episodic and contextual memory. In this framework, the common estimate of 200,000 EC layer II neurons projecting to the DG seemed adequate to convey such wealth of cortical information into the hippocampus and is comparable to other major hippocampal fields as CA3, CA1 or the subiculum that further elaborate on that information. However, our analysis reports that only about 80,000 neurons are responsible for delivering all cortical information to the DG (and CA3 and CA2). Furthermore, since MEC and LEC deal with different types of information and have segregated projections to the entire DG, it can be assumed that the 35,000 Reln+ neurons in MEC will be responsible for all spatial information delivered to the DG, and the 45,000 from LEC will convey all the contextual information. These populations seem small in the context of the hippocampal formation, as they represent about 4% of the 1 million GCs and about 10-20% of the 210,000-350,000 principal cells in other hippocampal fields like CA3, CA1 and subiculum (Arellano and Rakic, 2026) that also work with that information. Intuitively, one would expect a large pool of EC neurons allowing multiple combinatorial ensembles to convey the rich and complex contextual information necessary for hippocampal function to the DG. In turn, the relatively small number of EC neurons suggests that, beyond activation of different layer II neuron ensembles, signal and frequency encoding might play an important role conveying the complex contextual information expected to reach the DG.

### The perforant path shows richer and more extensive connectivity with GCs than previously considered

The updated number of dendritic spines in GCs suggests they receive about 10,000 excitatory synapses, 8,000 of them from EC axons. This estimate represents about double the number of excitatory synapses previously described (∼4,000; range 3,600-5,600) (Squire et al., 1989; Amaral et al., 1990; Patton and McNaughton, 1995; Derrick, 2007). The reason for such difference is that previous estimates were based on the number of GC dendritic spines calculated from estimates of total dendritic length and spine density using light microscopy. Although those spine density estimates are useful to compare between species or experimental conditions, it might not be accurate to describe number of spines, as it is prone to underestimation due to resolution limitations and the visual obstruction created by the dendrite and other spines (Feldman and Peters, 1979). For that reason, we used estimates of dendritic spine density based on dendrite reconstruction from EM serial sections, which warrants much higher accuracy detecting spines but has the downside of limited sampling. This approach has shown densities of ∼3 spines per micron across ML layers (Harris et al., 2022), that combined with the average GC dendritic length of ∼3,200 um (Desmond and Levy, 1982; Caceres and Steward, 1983; Claiborne et al., 1990; Cannon et al., 1999; Pierce et al., 2011; Buckmaster, 2012; Cole et al., 2020) provides an estimate of ∼10,000 spines per GC, about double of previous estimates (Amaral et al., 1990). This figure is supported by stereological studies reporting very similar accounts of ∼11,000 synapses per GC (West and Andersen, 1980; Cardoso et al., 2008a; Thind et al., 2010) and thus we are confident it might accurately reflect the total number of excitatory synapses on GCs. Out of those, about 75% or 8,000 are in the MML and OML (Crain et al., 1973; Geinisman et al., 1991, 1992; Cardoso et al., 2008b; Thind et al., 2010); present data), an estimate that is about double the traditional figure (∼4,000) of EC synaptic input to GCs mentioned above.

Such large number of contacts established by EC axons imply large synaptic divergence and suggests extraordinarily dense packing of synapses in the DG molecular layer, a possibility corroborated by EM studies showing that about 75% of the axonal varicosities in the MML and OML are multi-synaptic boutons that synapse with more than one spine, and in some cases up to 10 spines in mice (Popov and Stewart, 2009; Bosch et al., 2015). Those axons also exhibit some additional contacts in dendritic shafts that represent about 2% of the total excitatory synapses (Matthews et al., 1976; Thind et al., 2010; Bromer et al., 2018). Overall, this multi-synaptic approach seems an efficient way to maximize connectivity in the DG molecular layer.

### The perforant pathway is much more divergent than currently assumed

In connection with the point raised above, the relatively small number of EC neurons projecting to the DG reveals higher divergence in the EC-DG projection than previously assumed. reaching 1:30 and 1:20 for MEC and LEC respectively, about 6-fold and 4-fold the divergence of 1:5 described in currently models (Amaral et al., 1990; O’Reilly and McClelland, 1994; Patton and McNaughton, 1995; Becker, 2005; Aimone et al., 2006, 2009; Leutgeb et al., 2007; Newman and Hasselmo, 2014; Chavlis et al., 2017; Cayco-Gajic and Silver, 2019; Borzello et al., 2023).

At the synaptic level, the divergence is also remarkably high, as EC neurons are estimated to establish 90,000-115,000 synapses with GC. After considering a redundancy of 25% estimated by the model, each EC neuron might contact 68,000-86,000 neurons (for LEC and MEC, respectively) that represent 7-9% of the GC population. However, it is likely that the redundancy of the connectivity might be higher. Indeed, the 25% estimated here derives from EM studies reporting axons synapsing with two spines from the same dendrite or from different dendrites of the same neuron. However, given the small volume normally analyzed at the EM level, the method is not well suited to detect axons connecting with two distant dendrites of the same neuron, and therefore that 25% may represent an underestimation of the actual redundancy.

The estimate of 90,000-115,000 synapses per EC neuron is about 5-6 times higher than previous estimates of ∼18,000 synapses per EC neuron (Amaral et al., 1990; Patton and McNaughton, 1995), and is also about double to triple the number of synapses estimated for principal neurons from other hippocampal fields such as CA3 (∼40,000 synapses (Sik et al., 1993; Wittner et al., 2007)). This large number of synapses and connectivity spread matches morphological descriptions of widespread distribution of entorhinal axons in the molecular layer (Tamamaki and Nojyo, 1993) and suggests the scheme of connectivity leans towards maximizing GC sampling of EC input, contrary to the ultra-low sampling described in traditional models of pattern separation (Marr, 1969; Albus, 1971)(see functional implications of the updated model; e.g.: pattern separation).

### Surprisingly similar number of synapses on GCs from MEC and LEC

Both dendritic length and spine density in the MML and OML are remarkably similar, producing almost identical estimates of number of dendritic spines in those compartments. This is surprising, considering that LEC contains almost 30% more neurons than MEC projecting to GCs. This also implies that MEC neurons establish about 30% more synapses with GCs: 115,000 from MEC vs 90,000 from LEC. A potential reason to explain that difference could be that, since the MML might also receive input from hilar mossy cells, those synapses from mossy cells take certain percentage of the GC synapses, and the total synapses from MEC are closer to 90,000 from LEC. However, although speculative, our estimate of synapses from mossy cells into the MML suggests that they might represent a minimal part of the total, ∼40-160 synapses per GC, and therefore it seems that most of those estimated 115,000 synapses indeed originate from MEC.

Considering MEC input is closer to the soma of GCs than LEC input, an initial assumption could be that MEC input might have a stronger effect to activate GCs than LEC input. This, combined with the fact that MEC neurons establish 30% more synapses than LEC neurons, suggests that MEC activity might have stronger effect depolarizing GCs than LEC activity. However, this scheme is expected to be more complex, for example factoring feed forward inhibition by EC input, which can be differentially affected by MEC or LEC input. In fact, despite the rich entorhinal input, GCs show sparse activation that normally involves only 1-8% of the total GC population (Chawla et al., 2005; Ramírez-Amaya et al., 2005; Kee et al., 2007; Stone et al., 2011; Hainmueller and Bartos, 2018). Granule cells are intrinsically difficult to activate, a feature that has been related to their highly hyperpolarized resting potential, strong spike frequency adaptation and strong inhibitory input (Borzello et al., 2023).

### Mossy cell projections to the MML

The recent description of prolific projections from septal mossy cell to the MML in mice is intriguing, as no significant axonal arborization from mossy cells has been described in previous studies in rats. We speculate that ∼5,000 septal mossy cells from each hemisphere might establish a total of 4,000-16,000 synapses om the MML. Previous studies using *P. vulgaris* leucoagglutinin anterograde labeling of neurons in the hilus reported a diffuse axon plexus in the ipsilateral outer molecular layer (Amaral and Witter, 1989; Deller et al., 1994, 1995) around the level of the injection. And (Deller et al., 1995) also described projections to the MML in the contralateral side of the injection, but only at the same septo-temporal level. Furthermore, in contrast with the recent reports describing projections only from septal mossy cells, Deller and coworkers reported projections to the outer molecular layer from both septal and temporal injections, that were interpreted as originating from hilar GABAergic interneurons, likely expressing somatostatin or neuropeptide Y (Amaral and Witter, 1989; Deller et al., 1994, 1995). Another possibility is that this projection is present in mice but is absent in rats. Nonetheless, further studies, particularly at the ultrastructural level, are necessary to clarify the origin and extent of those projections and confirm if they target preferentially GCs, GABAergic interneurons or maybe a combination of both.

### Are there differences between male and female hippocampal connectivity?

Hippocampal studies have favored studying males to avoid the influence of cycling hormones present in females, as the hippocampus is sensitive to hormonal fluctuations (Gould et al., 1990; Zhang et al., 2008). Indeed, hormonal differences might be involved in the sexual dimorphism described in this structure: in rats the male hippocampus is larger, heavier and contains more neurons than the female hippocampus (Madeira et al., 1992; Andrade et al., 2000; Nuñez et al., 2003b). Sex dimorphism in the hippocampal formation is in agreement with the dimorphism in brain and body weight described in rats (Kakolewski et al., 1968; Goodrick, 1980; Madeira et al., 1988, 1991; Nuñez et al., 2003a, 2003b), mice (Wimer et al., 1988; Wimer and Wimer, 1989) and in guinea pigs (Severi et al., 2005), indicating this might be a general feature in rodents, and maybe in other species.

Importantly, it is not known if these differences are proportional across the hippocampus and males have a scaled-up version of the female hippocampus, or differences are uneven across fields and males and females might have different population and connectivity ratios. For the model, to avoid potential bias derived from mixing data from both sexes, we tried to be consistent using only data from male rats. However, there are cases like the neuronal populations of the EC, where detailed studies on layer II reporting separately the population of MEC and LEC are only available in females, while male data is either described for layer II or is described for MEC and LEC but for combined layers II and III. To estimate the population of MEC and LEC layer II for males, we used the average of 116,000 neurons in layer II, segregated using the 60:40 MEC to LEC ratio observed in females that is also similar to the 65:35 MEC to LEC ratio for combined layers II-III reported in one study in males (Arellano and Rakic, 2026). We obtained 70,000 neurons in MEC and 46,000 in LEC in males, compared to the 60,000 and 40,000 obtained for those fields in females (Arellano and Rakic, 2026). It must be noted that in our previous analysis, we estimated layer II MEC and LEC neuronal populations in males using a different criterion, that led to estimates of 76,000 neurons in MEC and 40,000 in LEC (Arellano and Rakic, 2026). However, those estimates derived from one single study, and we think they are less reliable than the ones reported here that are derived from the average of four studies.

Although the number of studies is small, it seems that males exhibit about 15% more neurons than females in the EC, a difference that is very similar to the 14% difference between sexes described for GCs in adult rats (Arellano and Rakic, 2026). It should be noted that the methodology used in some of those studies on GCs may not be optimal and therefore their results in absolute terms might not be accurate (see (Arellano and Rakic, 2026) for details). However, their ratios between sexes might be reliable, as the potential bias or error introduced is expected to influence both estimates similarly. Overall, these data suggest that the differences in neuronal populations across sexes might be proportional across fields, and the ratio of connectivity of the perforant path might be similar across sexes. However, other factors could play a role. For example, the smaller volume of the female ML might reduce GC dendrite length, potentially reducing the number of spines and thus excitatory inputs from the EC. In turn, spine density in females could be higher than in males and compensate for shorter dendrites, resulting in overall similar number of synapses in both sexes. Future studies might clarify which of those possibilities is correct.

### Functional implications of the updated model. E.g.: pattern separation

The increased divergence in the perforant path projection to the DG immediately brings to mind pattern separation, a function traditionally associated with the DG that allows disambiguation of two similar incoming input ensembles into two distinct patterns of output activity (Treves and Rolls, 1994; Hasselmo and Wyble, 1997). Pattern separation works through sparse encoding, meaning output is encoded by a small subset of neurons, reducing overlap between similar input activity patterns and maximizing coding capacity (Jung and McNaughton, 1993). Pattern separation was originally formulated in the cerebellum by (Marr, 1969; Albus, 1971) based on three distinct features of the mossy fiber projection to cerebellar granule cells: large divergence of the neuronal populations, sparse connectivity, and sparse output response mediated by widespread feed-forward inhibition (Marr, 1969; Albus, 1971; Cayco-Gajic and Silver, 2019). This functional model has also been proposed for other systems like the olfactory circuit of insects, as similar features are observed between olfactory projection neurons in the antennal lobe projecting to Kenyon cells in the mushroom bodies (Laurent, 2002; Perez-Orive et al., 2002; Cayco-Gajic and Silver, 2019) and for the hippocampus, based on the divergence of the EC projection to the DG and the sparse activity of GCs, with only about 1-8% active under normal stimulation (Chawla et al., 2005; Leutgeb et al., 2007; Alme et al., 2010; Danielson et al., 2016, 2017; GoodSmith et al., 2017).

In this context, the relatively low divergence, according to the traditional estimate of 1:5, of the EC to DG projection provided mild support for the functional model compared with the 1:600 divergence recorded in the mossy fiber to GC projection of the cerebellum that inspired the model (Eccles et al., 1967; Marr, 1969; Albus, 1971). Our updated model, however, describes a divergence of 1:20-30, one order of magnitude higher and similar to the divergence of 1:20-60 reported between olfactory projection neurons and Kenyon cells in Drosophila (Perez-Orive et al., 2002; Keene and Waddell, 2007).

The third characteristic of the Marr-Albus model, the sparsity of connectivity, is, however, not met in the EC-DG projection. Sparse connectivity in this context refers to the ultra-low synaptic convergence of source neurons into target cells. In the cerebellum, the convergence is 4:1, meaning each GC neuron receives input from an average of ∼4 mossy fibers, while in the mushroom body of Drosophila, the convergence of is 10:1, as Kenyon cells receive synapses from about 10 olfactory projection neurons (Eccles et al., 1967; Turner et al., 2008; Nguyen et al., 2023). In stark contrast, our model shows that the convergence of the EC projection to the DG for each of the subdivisions of the EC is 2-3 orders of magnitude higher: 3000:1, raising questions about the applicability of the Marr-Albus model to the perforant pathway. Nonetheless, a variety of experimental studies have described some form of pattern separation in the DG (Leutgeb et al., 2007; McHugh et al., 2007; Yassa and Stark, 2011; Neunuebel and Knierim, 2014), and therefore it might be achieved through a different computational scheme. In that regard, we hope our data might provide a reliable source for modelling this and other potential functions of the DG.

### Data-based modelling and speculative modelling

As mentioned in the introduction, there is a large amount of available information about the neuronal populations and synaptic connectivity of the hippocampal formation of the rat, mostly from males. This plethora of data is especially useful to build a complete and detailed model, and in some cases allows comparing data obtained from different labs or with different methods to certify their accuracy. That is the case, for example, of the neuronal populations in the EC of females, that are particularly consistent across studies (Arellano and Rakic, 2026) or the estimates of excitatory synapses on GCs, that are similar using stereology or calculating the number of dendritic spines. In that regard, we are confident some propositions of the model are pretty accurate. However, there are other features like the number of EC layer II neurons in MEC and LEC in male rats, or the number of Reln+ neurons in those fields that are approximations based on incomplete data or data from other species like mice and might be revised when more accurate data are available.

There is a third category of modelling that is based on speculation, for example when we estimate the potential synaptic input of mossy cells in the MML. In that case we can approximate the proportion of mossy cells potentially projecting to the MML, but there is no data regarding their target or the number of synapses they might establish. To fill the blank, we propose speculative values that nonetheless help gauging the extent of the projection and are, to certain extent, self-correcting. The initial proposal of 10-20% of the MML synapses originating from mossy cells seems unlikely when it is expressed as synapses per mossy cell, and therefore we propose a much lower value of 1-4% that matches previous reports and seems much more realistic. Thus, speculative modelling has an intrinsic value to propose testable hypothesis that can be validated logically or experimentally and helps putting findings in perspective for further discussion.

As quoted at the beginning of the manuscript, all models are wrong, but some are useful. And we hope this model, even if it might be wrong in some specific respects, will be useful to help understand information processing in the hippocampal formation and build more accurate models of hippocampal function.

## Conclusion

The model of the perforant pathway presented here provides a relevant updated review of quantitative data on the connectivity of the EC and the dentate gyrus, offering new insights: segregated cortical input for spatial and non-spatial contextual information reaches the DG through two small subpopulation of neurons in EC layer II that produce extensive synaptic coverage in the dentate gyrus, targeting a much larger subset of GCs, implying relatively large divergence. Overall, we think the updated model makes two important contributions: it provides detailed and accurate data to build hippocampal functional models and to establish comparisons with other species, and provides a starting point to study and discuss some of the predictions, that uncover new, interesting and potentially functionally relevant aspects of the flow of information in the hippocampal formation.

